# Enhanced nociceptive behavior and expansion of associated primary afferents in a rabbit model of cerebral palsy

**DOI:** 10.1101/2021.09.28.462176

**Authors:** E. J. Reedich, L. T. Genry, M. A. Singer, C. F. Cavarsan, E. Mena Avila, D. M. Boudreau, M. C. Brennan, A. M. Garrett, L. Dowaliby, M. R. Detloff, K. A. Quinlan

## Abstract

Spastic cerebral palsy (CP) is a movement disorder marked by hypertonia and hyperreflexia, and the most prevalent comorbidity is pain. Since spinal nociceptive afferents contribute to both the sensation of painful stimuli as well as reflex circuits involved in movement, we investigated the relationship between prenatal hypoxia-ischemia (HI) injury which can cause CP, and possible changes in spinal nociceptive circuitry. To do this, we examined nociceptive afferents and mechanical and thermal sensitivity of New Zealand White rabbit kits after prenatal HI or a sham surgical procedure. As described previously, a range of motor deficits similar to spastic CP was observed in kits born naturally after HI (40 minutes at ∼70-80% gestation). We found that HI caused an expansion of peptidergic afferents (marked by expression of calcitonin gene-related peptide; CGRP) in both the superficial and deep dorsal horn at postnatal day (P)5. Non-peptidergic nociceptive afferent arborization (labeled by isolectin B4; IB4) was unaltered in HI kits but overlap of the two populations (peptidergic and non-peptidergic nociceptors) was increased by HI. Density of glial fibrillary acidic protein (GFAP) was unchanged within spinal white matter regions important in nociceptive transmission at P5. We found that mechanical and thermal nociception was enhanced in HI kits even in the absence of motor deficits. These findings suggest that prenatal HI injury impacts spinal sensory pathways in addition to the more well-established disruptions to descending motor circuits. In conclusion, changes to spinal nociceptive circuitry could disrupt spinal reflexes and contribute to pain experienced by individuals with CP.

**Significance Statement:** Perinatal injuries that cause cerebral palsy (CP) typically involve global insults to the central nervous system and are capable of modulating development of both motor and sensory systems. Most individuals with CP experience pain, yet whether nociception is enhanced in this disorder is unexplored. Here, we demonstrate altered topographic distribution of nociceptive afferents in the spinal cord dorsal horn of neonatal rabbits that experienced hypoxic-ischemic (HI) injury in utero; these anatomical changes were associated with nocifensive behavior indicative of pain-like behaviors. Our findings suggest that CP-causative injuries alter spinal sensory pathways (not only descending motor circuits), contributing to increased pain in CP.

## Introduction

Cerebral palsy (CP) is the most common motor disability in children, affecting 1:500 live births (Oskoui et al., 2013). Chronic pain is the most prevalent co-morbidity of CP (McDowell et al., 2017; Novak et al., 2012): up to 77% of children with CP experience pain (McKinnon et al., 2020; Mckinnon et al., 2018). Individuals with CP experience chronic pain at significantly higher rates than the rest of the population (Badia et al., 2014; Doralp and Bartlett, 2010; García Jalón et al., 2021; Westbom et al., 2017). Pain decreases quality of life (Findlay et al., 2016), yet the pathophysiology of pain is rarely researched in the context of CP. This is likely due to the pervasive view that pain in individuals with CP is a symptom arising from secondary effects of physical stressors, including spasticity, hip subluxation, and joint contractures. Individuals with severe CP (as measured by Gross Motor Function Classification System (GMFCS) V) do experience more pain (García Jalón et al., 2021; Mckinnon et al., 2018); however, CP-affected individuals across all GMFCS levels report pain (Flanigan et al., 2020; Sienko, 2018). Even individuals born very prematurely (<26 weeks gestion), who do not develop CP, report recurrent pain and altered thermal sensitivity as adults (Walker et al., 2018). Thus, we hypothesized that the same trauma that can impact developing corticospinal tracts in CP could impact development of nociceptive circuitry in the spinal cord.

There is a wealth of evidence regarding the interrelatedness of sensory and motor performance (Cooper et al., 1995; Gupta et al., 2017; Matusz et al., 2018; Zarkou et al., 2020). Somatosensory deficits are present in CP, and their presence could contribute to motor dysfunction. Typically, primary afferent input onto spinal circuitry is a key mediator of functional motor performance. Proprioceptive and tactile afferent feedback informs the central nervous system (CNS) of motor command outcomes and sculpts ongoing drive to motoneurons. In particular, the role of primary nociceptive afferent fibers in motor function is critically overlooked—especially after CNS damage. Importantly, there are a few exceptions, including the restriction of motor unit activation in response to excessive stretch, detection of dangerous pH and other metabolite levels during muscle fatigue, and the exercise pressor reflex and motor unit modulation during exercise-induced fatigue (Laurin et al., 2015; Zając et al., 2015).

Cutaneous nociceptors (mechanosensitive C, thermosensitive C and Aδ neurons) are small, pseudounipolar neurons in the dorsal root ganglion (DRG) that are analogous to Type III (Aδ) and Type IV (C) primary afferent fibers from the muscle, though their axons terminate in different dorsal horn laminae (Kandel et al., 2013; Lallemend and Ernfors, 2012; Laurin et al., 2015). Importantly, nociceptive and pain information can be relayed from muscle and joints via the nociceptive primary afferent reflex arcs, modulating motor output by generating a withdrawal reflex (Kandel et al., 2013; Kimura et al., 2004). The interconnection between motor and sensory systems can make it challenging to parse out their relative contribution to functional impairments.

As in other CNS disorders, there is a strong correlation between the integrity of the somatosensory system (including primary afferent input) and motor performance in CP (Gupta et al., 2017; Zarkou et al., 2020). Tractography and physiology experiments in individuals with unilateral CP revealed that integrity of somatosensory connections has a relatively larger impact on hand function compared to the integrity of motor connections (Gupta et al., 2017), and stimulation of peripheral nerves, which provides additional sensory feedback, improved motor performance in individuals with diplegic CP (Bumin and Kayihan, 2001; Kaelin-Lang, 2008).

Physical insults during pregnancy have been linked to later development of autism, schizophrenia, and the motor impairments that are hallmarks of CP (Marriott et al., 2017), but it is not known if these prenatal events impact nociceptive afferents. Neural development can be negatively impacted by relatively common conditions in the perinatal period, including maternal infection and inflammation, placental insufficiency, and difficult birth. These conditions can all result in varying degrees of hypoxia-ischemia (HI) and injury to the developing CNS. Effects of prenatal HI include white matter injury, cortical damage and motor deficits (Buser et al., 2010; Derrick et al., 2004; Drobyshevsky and Quinlan, 2017; Rocha-Ferreira and Hristova, 2016), but the effect on nociceptors is completely unstudied. Notably, cell death in the deep dorsal horn (where synaptic targets of nociceptors are located) is present after HI in rabbits and in cases of infant death after hypoxia (Clancy et al., 1989; Drobyshevsky and Quinlan, 2017; Sladky and Rorke, 1986). Dorsal root ganglia (DRG) neurons undergo perinatal growth and refinement of projections during the period of susceptibility to injuries causing CP (Marmigère and Ernfors, 2007; Olson and Luo, 2018), though afferent plasticity is retained in adulthood (Jiang et al., 2016; Tan et al., 2012). Changes to nociceptive DRG neurons are entirely unexplored in the context of CP.

In this study, we have chosen to focus on nociceptors at the spinal level using the rabbit model of CP. Investigating the relationship between motor and sensory dysfunction in CP requires using an animal model that displays both phenotypes of motor dysfunction and enhanced nociception. Unfortunately, rodents, the most commonly used models, can sustain very severe brain injuries yet display mild or nonexistent motor deficits (Fan et al., 2005; Rice et al., 1981)HI kits show pronounced motor dysfunction (Cavarsan et al., 2019; Derrick et al., 2007, 2004; Shi et al., 2021). Furthermore, emerging evidence suggests that nociceptors may link afferents to hyperreflexia. Nociceptors impact movement by modulating spinal reflex circuits (Black, 2019; Lucas-Osma et al., 2019), and nociceptor input to reflex circuits is a feasible potential contributor to disrupted reflex responses (Hari et al., 2021) particularly in those with CP who may suffer from chronic pain. Monosynaptic stretch reflexes can be potently suppressed via nociceptor inhibition (Black, 2019; Lucas-Osma et al., 2019). In the case of people with CP, rather than suppression of the monosynaptic stretch reflexes, over active nociceptors may drive hyperreflexia and reflex irradiation (Achache et al., 2010; Leonard et al., 1991; Leonard and Hirschfeld, 1995). Whether this mechanism is contributing to spasticity and hyperreflexia in CP is the focus of ongoing study. Thus, not only could expanded distribution of nociceptive afferents contribute to the perception of pain by higher brain centers, but also to hyperreflexia and disruptions in motor coordination that are the hallmarks of CP.

Thus, we examined nociceptors and altered nociception at the level of the spinal cord in rabbit kits that were subjected to HI injury in utero (40 minutes at ∼70-80% gestation) using both immunohistochemistry to quantify molecular markers for nociceptors and in vivo behavioral testing at an age roughly corresponding to human toddlers (postnatal day [P] 5). We found that rabbit kits exposed to prenatal HI demonstrated increased nociceptive behavior in response to cutaneous mechanical and thermal stimuli. This increased nociceptive response corresponded to aberrant distribution and density of nociceptive primary afferent fibers but not astrogliosis at this age. Interestingly, our data show that enhanced forepaw and hindpaw sensitivity occurs independent of motor deficits produced by this model. Together, these results indicate that sensory circuits are vulnerable to perturbation by perinatal HI injury, and suggest that neuropathic pain could accompany motor deficits characteristic of CP.

## Materials and Methods

### Animal model

Experiments were performed in accordance with the ARRIVE guidelines and the United States National Institutes of Health’s Guide for Care and Use of Laboratory Animals. Approval of the University of Rhode Island’s Institutional Animal Care and Use Committee was obtained for all experiments performed in this study. 66 kits of both sexes from 15 different litters originating from 9 dams were used for this study; some dams provided more than one litter of rabbits. The inherent variability in behavioral data warranted large sample sizes. All efforts were made to minimize animal suffering and to reduce the number of animals used.

### Hypoxia-ischemia surgery

Pregnant New Zealand White rabbits (bred in house or ordered from Charles River Laboratories, Inc., Wilmington MA), underwent HI procedures as previously described (Derrick et al., 2004). Briefly, at ∼70-80% gestation (day 22 -26 of 31.5 days typical gestation period), dams were premedicated with ketamine and xylazine (25 mg/kg and 3.75 mg/kg, respectively), anesthetized with isoflurane (1-3%; inhaled via V gel tube) and occasionally treated with buprenorphine (0.01 - 0.24 mg/kg, as needed). Indications for treatment with buprenorphine include elevated heart rate (over 220BPM); prolonged rapid, shallow breathing / panting; elevated CO_2_ levels (due to panting). The left or right femoral artery was isolated. A Fogarty balloon catheter (2-4 French) was inserted and advanced to the level of the bifurcation of the descending aorta, above the uterine arteries, and inflated to completely occlude blood flow for 40 minutes to cause HI in the fetuses. Placement of the catheter and confirmation that blood flow was occluded was performed with ultrasound. Dams subjected to sham surgical procedures underwent the same anesthesia protocols as the HI dams but without insertion and/or inflation of the catheter.

After the procedures, dams recovered and later gave birth to kits on the expected due date +/-two days. Sham and HI litters included in this study were all born at term (>92% of gestation). Neonatal kits of both sexes were characterized for motor deficits, nociceptive behavior, and used for anatomical investigations as described below. Since we examined behavior in neonatal animals far from sexual maturity (NZW rabbits typically reach sexual maturity at ∼6 months of age), and whose external genitalia were not yet visible, we grouped together male and female kits.

### Behavior

There is a wide range in motor phenotypes after HI, with some kits showing motor deficits while others appear unaffected. Categorization of motor deficits was performed using a modified Ashworth scale, observations/tests for activity, locomotion, posture, righting reflex, and muscle tone as previously described (Derrick et al., 2004). Based on these tests, kits were labeled as “motor-affected HI” or “motor-unaffected HI.” Motor-affected HI kits had modified Ashworth muscle tone scores of _≥_3 in one or more evaluated joints; Ashworth scores did not exceed 2 in any joints of motor-unaffected HI kits.

#### von Frey test for mechanical nociception

Neonatal rabbit kits of both sexes were tested for mechanical hypersensitivity using von Frey filaments. Animals were allowed to acclimatize in a chamber on an elevated gridded (8 mm x 8 mm) surface. von Frey filaments (Touch Test® Sensory Evaluators; range: 1-26 g) were applied in ascending order to the plantar surface of the forepaws and hindpaws, just proximal to the digits of the kit. See inset in Figure 1 which depicts location of stimulus application. If paw withdrawal occurred in response to application of the 1 g von Frey filament, filaments (0.6-0.008 g) were applied in descending order until no response was observed. Each filament was tested at least five consecutive times per paw to ensure the observed reaction was in response to the filament and not general movement of the animal. The force (in g) of the filament that elicited a consistent withdrawal response was recorded as the paw withdrawal threshold. If no response was elicited by application of the 26 g von Frey filament, that paw was recorded as non-responsive and given a score of 60 g (which is the subsequent filament). Filament forces eliciting withdrawal of left and right forepaws and hindpaws were entered independently for statistical analyses, since motor deficits are frequently disparate across limbs and responsiveness to stimuli was therefore not assumed to be equal across limbs.

**Figure 1:**
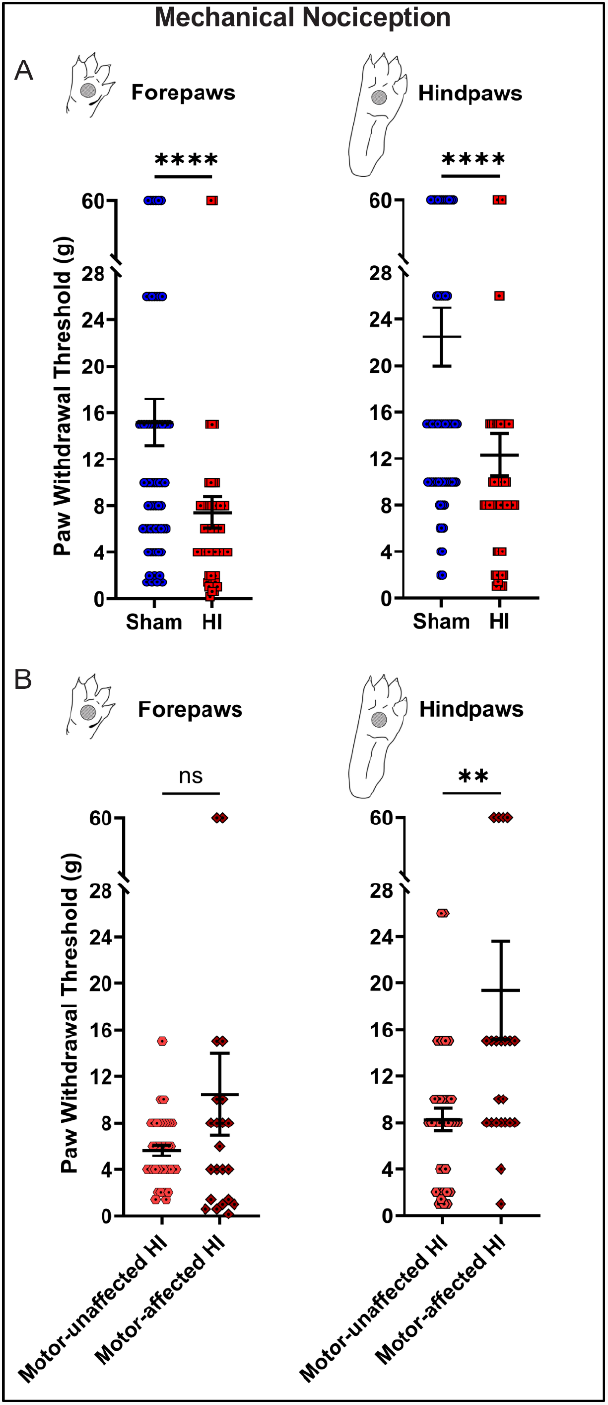
HI kits exhibit pain-like behavior in response to mechanical stimuli. **A:** Forepaw (left) and hindpaw (right) von Frey scores at P5. Sham-operated controls (blue) have higher scores, indicating less mechanosensitivity, while HI kits (red) display greater mechanosensitivity (lower scores). **B:** HI kits from (A) were separated by whether they show motor deficits (motor-affected) or no deficits (motor-unaffected). There was no difference in forepaw withdrawal threshold (left) but hindpaw withdrawal (right) scores were higher in motor-affected HI kits, potentially indicating less sensitivity or impaired ability to withdraw. Data are represented as mean ± SEM; N=64 sham and N=60 HI paws from N=32 sham and N=30 HI rabbits. **P<0.01, ****P <0.0001; Mann-Whitney tests.

#### Hargreaves’ test for thermal nociception

Thermal sensitivity was tested using a Hargreaves apparatus. Specifically, a model 336G Plantar/Tail Stimulator Analgesia Meter (IITC Life Sciences, Woodland Hills, CA) was used to apply a beam of radiant heat (4×6 mm area; 80% intensity; 8 s max duration) to the plantar surface of each hindpaw and forepaw. Rabbit kits were allowed to acclimatize in a chamber on a glass surface, and then the skin on the plantar surface of each paw proximal to the digits was tested three times with at least 30 seconds between applications. See insets in Figure 2 for schematic depicting the location of stimulus application to the forepaws and hindpaws. The beam of radiant heat increased temperature gradually from rest (20-30 °C) to a maximum of 83 °C at 8 s. Latency to paw withdrawal was recorded; if the animal did not withdraw its paw, that trial was recorded as a non-response and given a latency of 9 s. For each paw, the stimulus was applied 3 times, and the withdrawal latencies were averaged to generate a single score per paw. Since weight distribution and function are different between forelimbs and hindpaws, withdrawal latencies from the forepaws were examined separately from hindpaws. Each group contains individual data points generated from the left or right paws.

**Figure 2:**
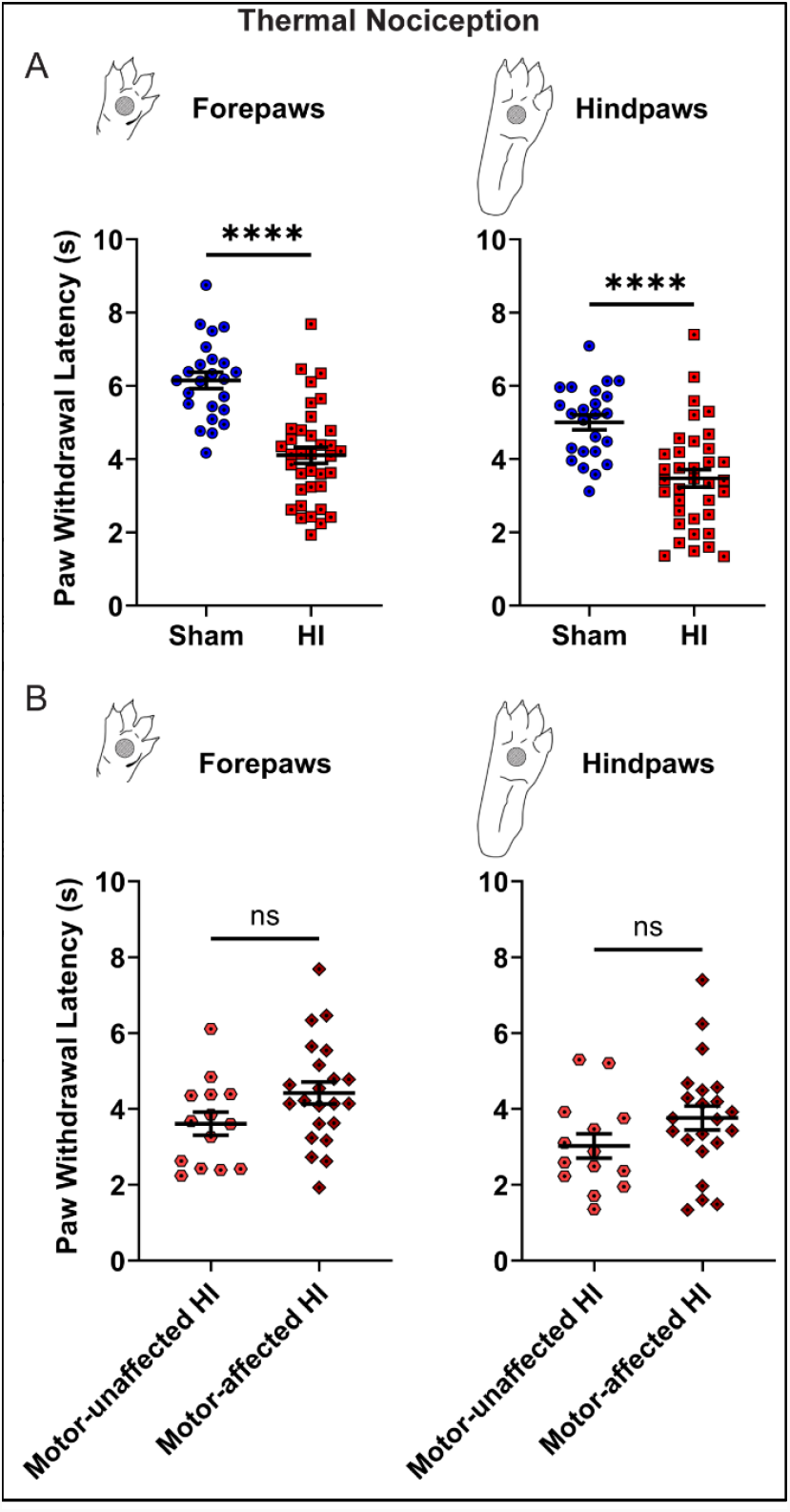
HI kits display pain-like behavior in response to thermal stimuli **A:** Forepaw (left) and hindpaw (right) withdrawal latencies during Hargreaves’ testing at P5. Shorter latencies to paw withdrawal in response to radiant heat are observed in HI kits compared to sham animals, indicative of enhanced thermal sensitivity. **B:** HI kits from (A) were separated by whether they show motor deficits (motor-affected) or no deficits (motor-unaffected). There was no difference in forepaw or hindpaw withdrawal scores in motor-affected HI kits. Data are represented as mean ± SEM; N= 24 sham and N=36 HI paws from N=12 sham and N=18 HI rabbits. ****P <0.0001; unpaired t-tests.

### Immunohistochemistry (IHC)

Kits were deeply anesthetized by intraperitoneal injection of ketamine/xylazine (100 mg/kg and 20 mg/kg, respectively) or sodium pentobarbital before tissue harvest at P5. Transcardial perfusion was performed using chilled 0.01 M phosphate buffered saline (PBS; pH 7.4) until tissue blanching was observed, then 4% paraformaldehyde (PFA) (in PBS) was infused at a constant flow rate of approximately 5-8 mL/min (depending on rabbit kit size). Perfused spinal cords were dissected and post-fixed in 4% paraformaldehyde for ∼24-72 hrs. Next, fixed spinal cords were washed with PBS and cryoprotected in 30% sucrose (in PBS) until sinking was observed. Cryoprotected spinal cords were embedded in optimal cutting temperature (OCT) compound and frozen at −80 °C until cryostat sectioning (Leica Lm 1850, Germany). Transverse lumbar spinal cord cryosections were collected in serial at 30-μm or 40-μm thickness; to avoid repeat regions of analysis, every 15th section was used for analysis. Manufacturer Information, catalog number and lot information for all immunohistochemical antibodies and lectins used are provided in Table 1.

**Table 1.**
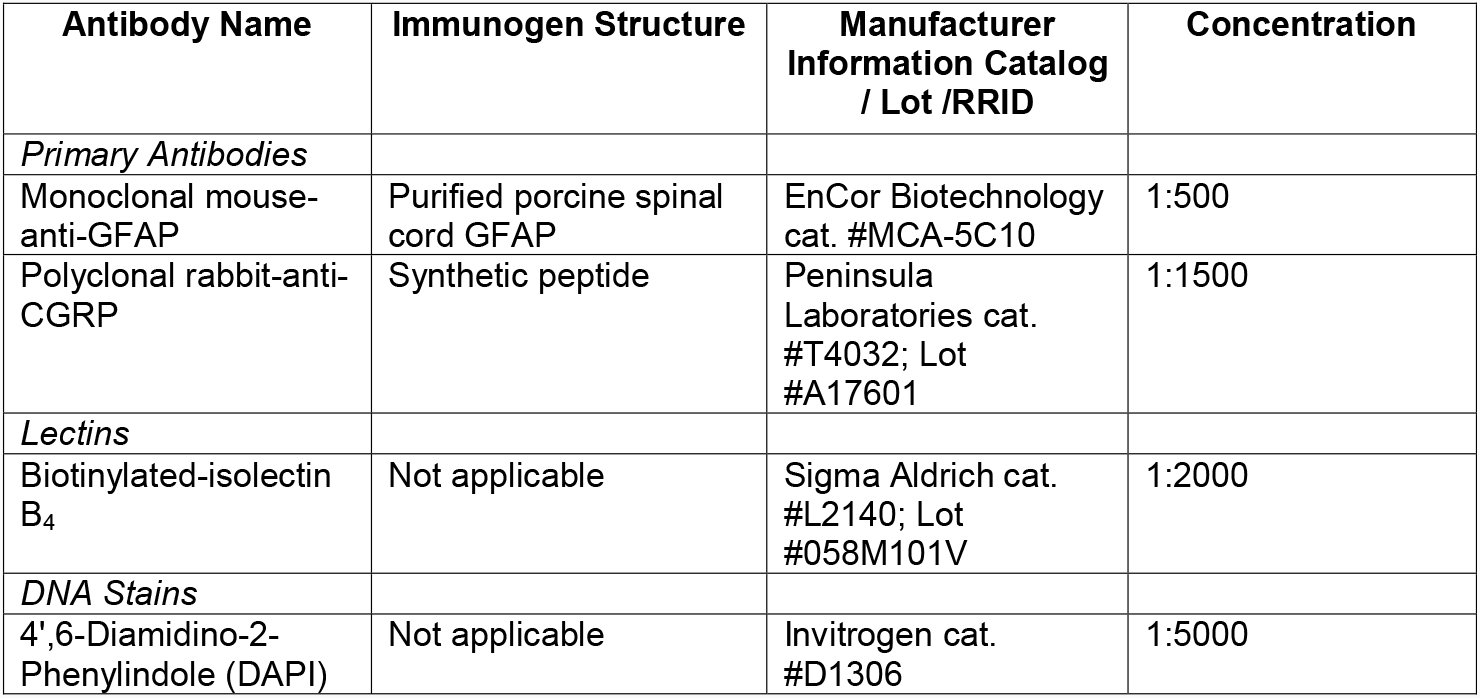
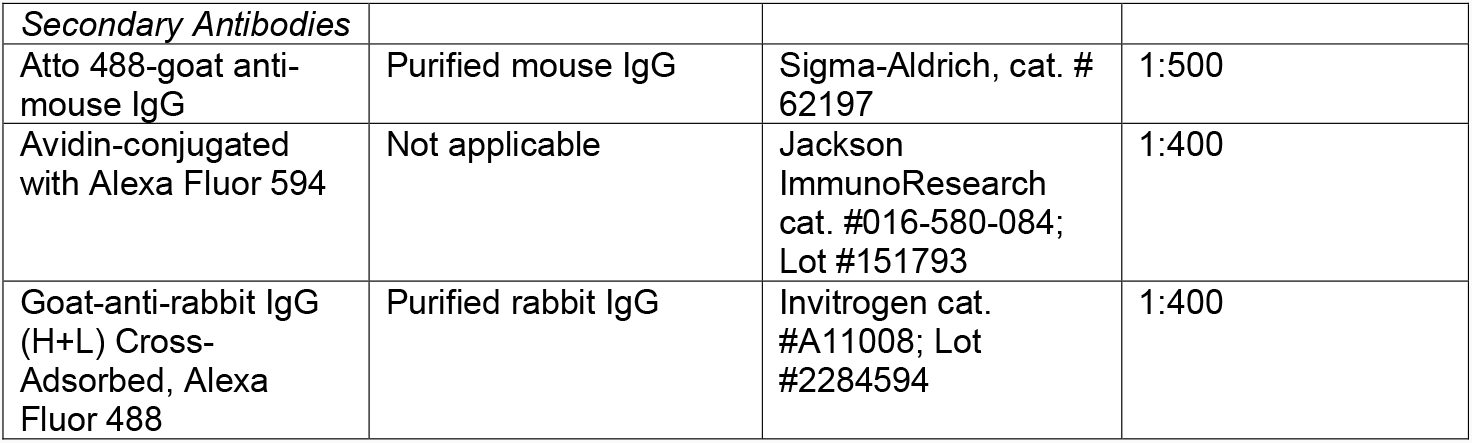
Immunohistochemical Reagents

#### IHC for glial fibrillary acidic protein (GFAP)

Slides were thawed, dried in an oven at 37 °C to improve section adherence, then rehydrated in PBS. Sections were blocked in 0.2% Triton X-100 and 5% normal horse serum (NHS) in PBS for 30 minutes at room temperature. Following blocking, slices were incubated in primary antibody solution containing mouse-anti GFAP antibody (diluted 1:500 in 0.1% Triton X-100 and 1% NHS in PBS;) overnight at room temperature. Sections were washed in PBS and then were incubated in secondary antibody solution consisting of Atto 488-goat anti-mouse IgG (whole molecule) (diluted 1:500 in 0.1% Triton X-100 and 1% NHS in PBS,) for 2 h at room temperature. Nuclei were labeled with 4’,6-Diamidino-2-Phenylindole (DAPI) and slides were mounted using Fluoromount mounting medium (SouthernBioTech). Z-stack photomicrographs of dorsal and ventral lower lumbar spinal cord were acquired at 10x magnification using a Nikon Eclipse Ti2 inverted confocal microscope. Maximum intensity projections of the green channel were generated and stitched (if possible) using Image Composite Editor (Microsoft). Grayscale images were imported into ImageJ and blinded densitometric analysis of GFAP immunofluorescent signal (mean gray value) in regions of interest containing dorsal columns, superficial dorsal gray matter, lateral spinothalamic tract, and ventral spinothalamic tract (divided by background fluorescent signal) was performed using Image J (NIH). Background fluorescent signal was the mean gray value of adjacent unlabeled gray matter. The grey or white matter in our regions of interest was targeted to encompass the entire region of interest but could also contain portions of other areas.

#### IHC for primary afferent fibers

Sections were rinsed in PBS, blocked in 10% normal goat serum and 0.2% Triton-X100 in PBS for 1 hour at room temperature followed by a 30-minute incubation with rabbit-to-rabbit blocking reagent (ScyTek Laboratories, Inc., catalog #: RTR008). Sections were then incubated with a solution of biotinylated IB4 (1:2000, Sigma) and anti-calcitonin gene-regulated peptide in 0.1% Triton X-100 in PBS at room temperature for 48 h. After three washes with PBS, sections were incubated in avidin-conjugated with Alexa Fluor 594 and goat-anti-rabbit Alexa Fluor 488 in 5% goat serum in PBS for 2 hours at room temperature. Following three washes, sections were rinsed in PBS and cover slipped with FluorSave Reagent (Calbiochem, catalog #: 345789, Bedford, MA). Three images of the left and right lumbar dorsal horn were taken for each rabbit at the same magnification, lens aperture and exposure time on a Leica DM5500B epifluorescent microscope and captured with Slidebook software. Images were averaged to calculate values per animal. The proportional area of IB4, CGRP and colocalized immunoreactive tissue within the cap of the dorsal horn (laminae I-III) was quantified in ImageJ as previously described (Detloff et al., 2016). The proportional area of CGRP+ fibers that expand in dorsal horn ventral to the IB4+ band was measured in the same sections.

### Statistical Analyses

All statistical analyses were performed using GraphPad Prism (GraphPad Software Inc.; San Diego, CA, USA). For each dataset, normality of distributions was tested using D’Agostino and Pearson tests. For normally distributed datasets with equal variances, unpaired t-tests were used to identify statistically significant differences between groups. For datasets in which one or more distributions was non-normally distributed, Mann-Whitney tests were used.

## Results

In this study, we investigated the link between prenatal HI injury and nociception in a rabbit model of CP. Rabbit kits exposed to prenatal HI and age-matched sham animals underwent testing for enhanced nociceptive reflexes in response to mechanical and thermal stimuli at P5 using von Frey and Hargreaves’ testing, respectively. After prenatal HI injury, neonatal rabbit kits displayed increased nociceptive behavior in response to mechanical and thermal stimuli that manifested even in the absence of motor deficits. HI kits demonstrated significantly greater sensitivity to von Frey filaments applied to the paws compared to sham kits, withdrawing paws in response to application of lower forces (forepaws, P<0.0001; hindpaws, P<0.0001) (**Figure 1**). Hindpaw von Frey scores of motor-unaffected HI kits (mean: 8.2 g;) were significantly lower than that of motor-affected HI kits (mean: 19.4 g; P=0.0087) while forepaw withdrawal thresholds were equivalent (P=0.9908).

Similar to von Frey, the Hargreaves’ test revealed paw hypersensitivity in HI kits. HI kits had reduced paw withdrawal latencies compared to sham rabbits in response to targeted radiant heat (Figure 2), indicating increased thermal sensitivity (forepaws: P<0.0001; hindpaws: P<0.0001). Paw withdrawal latencies to noxious thermal stimulation were comparable between motor-unaffected HI (forepaw latency: 3.6 s; hindpaw latency: 3.0 s) and motor-affected HI kits (forepaw latency: 4.4 s; hindpaw latency: 3.8 s) (P=0.0740 and 0.1279, respectively).

We hypothesized that mechanical and thermal hypersensitivity in HI rabbit kits may be caused by spinal astrogliosis and/or expansion of nociceptive afferents in the spinal cord dorsal horn. To determine whether prenatal HI leads to long-lasting postnatal astrogliosis in gray and white matter spinal cord regions involved in cutaneous sensation or nociception, the degree of glial fibrillary acidic protein (GFAP) expression in four lumbar spinal cord regions of interest (ROIs) was quantified. These ROIs included tracts/laminae involved in cutaneous sensation or nociceptive neurotransmission: the dorsal column, superficial dorsal gray matter, ventral spinothalamic tract, and lateral spinothalamic tract. Densitometric analysis revealed no significant differences in GFAP labeling in these ROIs at P5 (Figure 3). Baseline GFAP immunoreactivity in the white matter of neonatal rabbits found here is consistent with the nervous system expression profile established in mice (Yoon et al., 2017) and may reflect the role of astrocytes in myelination during early postnatal development (Landry et al., 1990; Liedtke et al., 1996; Nash et al., 2011). No evidence was found to support a contribution of neuroinflammation to increased nociception at this age.

**Figure 3:**
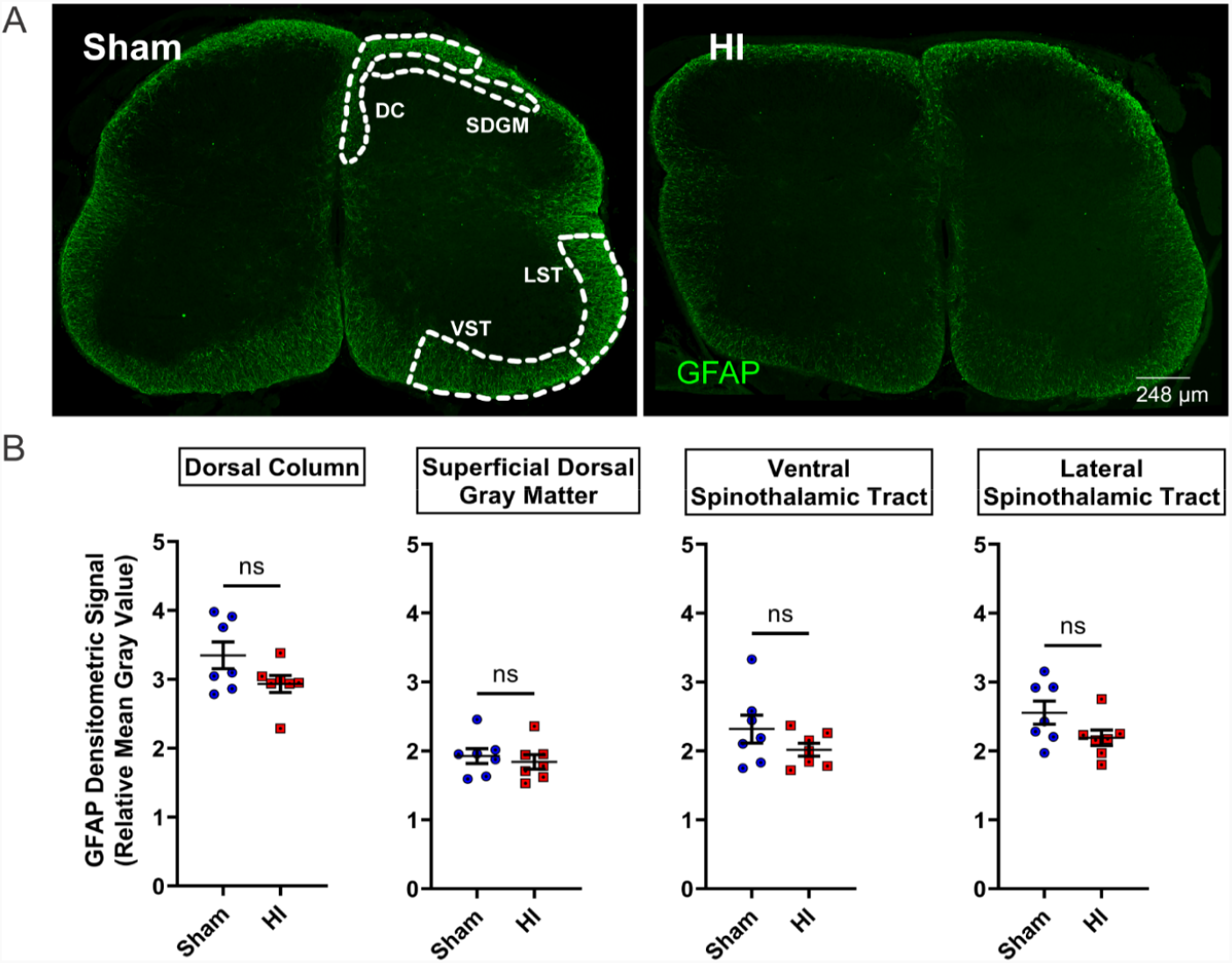
Astrogliosis is not present in HI rabbits at P5. **A:** Transverse lumbar spinal cord sections from sham (left) and HI (right) rabbits immunostained for GFAP (green). Scale bar applies to both images. **B:** Quantification of GFAP densitometry in spinal regions containing tracts/laminae of interest: dorsal column (DC), superficial dorsal gray matter (SDGM), ventral spinothalamic tract (VST), and lateral spinothalamic tract (LST). GFAP expression was unchanged in all ROIs in HI spinal cords (ns, not significant; unpaired t-tests). Data are represented as mean ± SEM; N=7 rabbits per group.

To determine whether prenatal HI affects primary afferent distribution and density within the dorsal horn of the spinal cord, lumbar spinal cords (which include hindpaw dermatomes) from sham and HI kits were labeled for CGRP and IB4 to identify peptidergic and non-peptidergic afferents, respectively. In other species, small diameter fibers emanating from the dorsal root ganglia that contain CGRP and bind IB4 transmit nociceptive information from the periphery to the superficial dorsal horn of the spinal cord (REF). Similar to the rodent, distributions of these two nociceptor subpopulations had little overlap in the dorsal horn of sham kits (**Figure 4 A-A’’**), however, HI led to an increased overlap in the topographic distribution of these two fiber types (**Figure 4 B-B’’**). In addition, HI kits exhibited increased CGRP-positive fibers in the deep dorsal horn (**Figure 4 C-C’’**) that were not present in sham kits. Quantification of the proportional area of each afferent subpopulation independently revealed that CGRP+ fibers (**Figure 3D, G**) but not IB4+ (**Figure 4E**) fibers exhibited increased density in the superficial and deep dorsal horn. Despite no increase in the density or distribution of IB4-positive fibers in the dorsal horn, there was significant colocalization of CGRP and IB4 labeling in the dorsal horn of HI kits compared to sham kits, suggesting that HI may induce plasticity or phenotype switching of these nociceptor populations.

**Figure 4.**
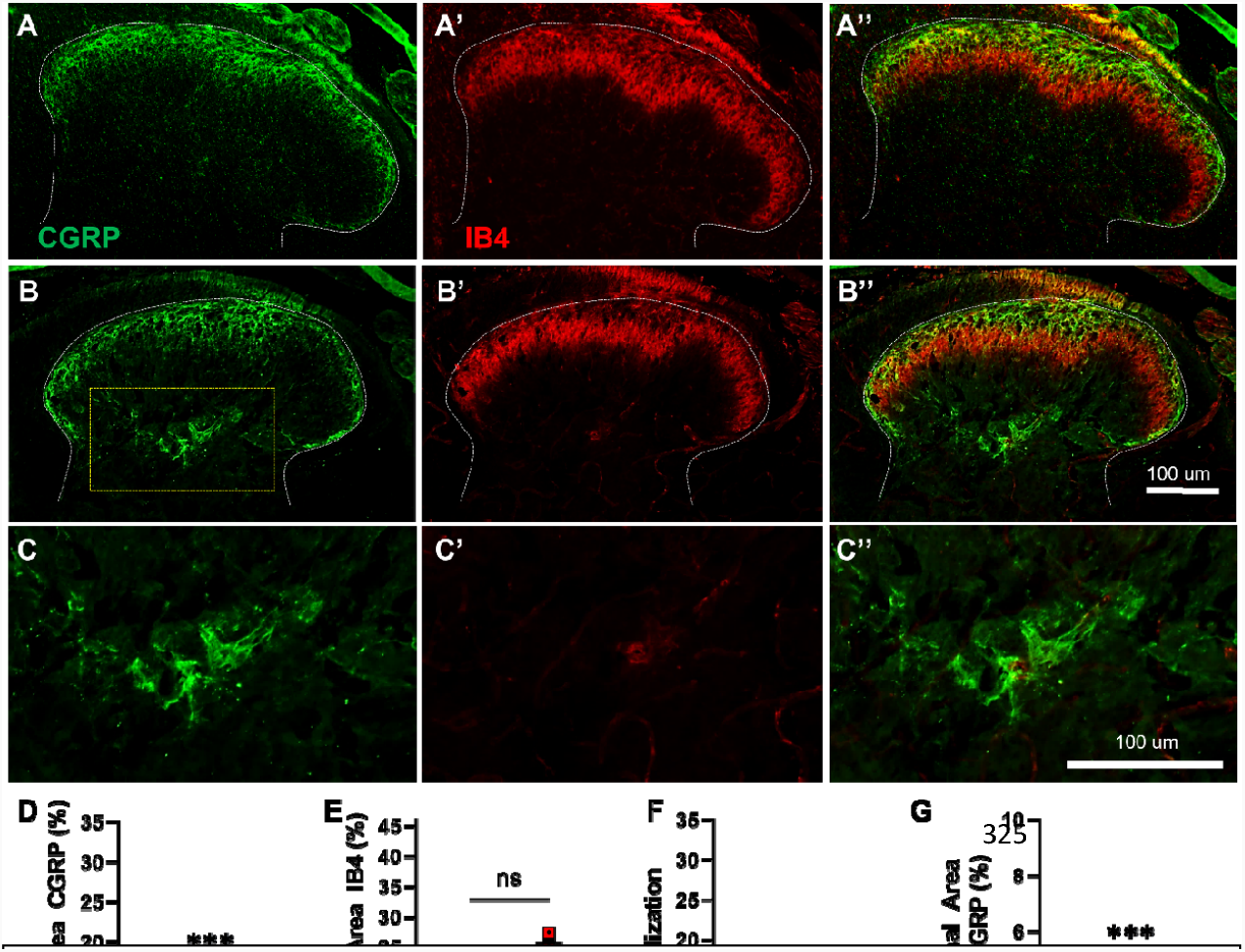
Increased density and overlap of nociceptive primary afferent fibers in HI kits at P5. Representative images of the lumbar dorsal horn of sham (**A-A’’**) and HI (**B-B’’**) kits immunolabeled for calcitonin gene-regulated peptide (CGRP; **A, B, C**) or labeled by isolectin B-4 (IB4; **A’, B’, C’**). Note the increased presence of CGRP-positive fibers in the deep dorsal horn in the HI but not the sham spinal cord (**C-C’’**). The proportional area of CGRP-positive tissue was significantly increased in the superficial and deep dorsal horn of HI rabbits compared to sham (**D, G**), while the density and distribution of IB4-positive afferents was unchanged between groups (**E**). The degree of overlap of these two nociceptor populations can be viewed in the merged images (**A’’, B’’**) and was significantly increased in HI kits compared to sham kits (**F**). Data are represented as mean ± SEM; individual data points represent each animal (n=5 sham and n=6 HI kits) from averages of 3 separate images of left and right dorsal horns per animal. **P<0.01, ***P<0.001; unpaired t-tests were used for statistical analysis.

## Discussion

Experimental animal models that mimic human disease with high fidelity are more likely to drive meaningful discovery of therapeutic interventions to treat such diseases. Here, we use a preclinical model that displays prominent motor deficits to show that nociceptive afferents are also disrupted, in addition to the previously characterized damage to the cortex, thinning of the corticospinal tract, loss of spinal motoneurons and altered motoneuron excitability (Buser et al., 2010; Drobyshevsky et al., 2007; Drobyshevsky and Quinlan, 2017; Steele et al., 2020). Importantly, both rabbits after HI and human infants after asphyxiation show cell death in the dorsal horn (Clancy et al., 1989; Drobyshevsky and Quinlan, 2017; Sladky and Rorke, 1986). Moreover, this is one of few studies that utilize quantitative assessments of nociception in the rabbit, despite the rabbit’s extensive use in biomedical research (Lei et al., 2017; Mans, 2020; Yoshioka et al., 1996). Our findings indicate that after prenatal HI, increased topographic distribution of peptidergic nociceptive primary afferent fibers in the dorsal horn is present along with a hypersensitive withdrawal response that is suggestive of allodynia.

There is substantial information regarding the formation of sensory connections during development in the rodent, cat and primate (Beggs et al., 2002; Brown and Fuchs, 1975; Fitzgerald and Jennings, 1999; Fitzgerald and Swett, 1983; Jennings and Fitzgerald, 1996; Ralston and Ralston, 1979; Scheibel and Scheibel, 1968; Swett and Woolf, 1985; Zhang et al., 1997), however, there is a paucity of information regarding primary afferent distribution in the developing or adult rabbit. This is the first study to show that nociceptors are altered in an animal model of CP and one of the first to examine primary afferent fiber distribution in the developing rabbit. The increased overlap and density of peptidergic nociceptive afferent fibers in the dorsal horn of HI kits may indicate intrinsic changes within nociceptive DRG neurons. In normal rodents, CGRP and IB4 are rarely co-expressed in primary sensory neurons of the DRG; these markers normally indicate distinctive subpopulations of nociceptors (Marmigère and Ernfors, 2007; Orozco et al., 2001). In chronic pathological situations, the distribution of the axonal arbors within the dorsal horn exhibits significant overlap, and there is some indication that the degree of coexpression of these two nociceptive markers increases (Detloff et al., 2016; Price and Flores, 2007; Sliwinski et al., 2018). Despite significantly lower von Frey scores in HI kits indicating increased mechanosensitivity, non-peptidergic IB4-positive mechanical nociceptors, which are activated by noxious mechanical stimuli, were not found to have expanded afferent arborizations in the dorsal horn. It could be that these IB4-positive nociceptors are hyperexcitable, and that altered activity, not anatomy, contributes to pain-like behavior in response to mechanical stimuli in HI kits. In other rodent models of nervous system injury, chronic pain and paw hypersensitivity is associated with dysfunctional nociceptors that exhibit robust anatomical and electrophysiological plasticity (Bedi et al., 2010; Detloff et al., 2016, 2014; Dougherty et al., 2004; Moy et al., 2018; Zhang et al., 2013); further, emerging evidence suggests that nociceptive primary afferent input impedes motor output following CNS injury (Keller et al., 2017, 2019, 2018). In a series of studies, Keller et al. demonstrated that nociceptor depletion reduces muscle contractures and spasticity of the hindlimbs and improves overall locomotor performance following spinal cord injury. Nociceptor silencing via activation of inhibitory 5HT_1D_ receptors has also been found to be helpful in reducing clonus and spasms after spinal cord injury (Lucas-Osma et al., 2019). Taken together, these findings indicate that altered activity in nociceptors contributes to motor deficits in cases of CNS injury. While the relationship with motor deficits is particularly important in CP, the methods used in this study are unable to distinguish CGRP and/or IB4 positive primary afferent fibers emanating from the skin, muscle or viscera, and behavioral assessments here only assess cutaneous afferent responses. Moreover, as dorsal root ganglia, where nociceptive cell bodies reside, were not systematically harvested in this study, we cannot determine if the altered topographic distribution of these afferents may represent a sprouting response or it may be indicative of a change in the population of DRG neurons which express these nociceptive markers. Populations of primary sensory neurons have been identified in many mammalian species from rodents to humans, and size, molecular markers and functions are mostly conserved (Shiers et al., 2020). While isolated anatomical reports suggest that the primary sensory neurons of rabbits may be anatomically similar to other mammals (Kusunoki et al., 1992; Mori, 1986), a systematic evaluation of the anatomical and physiological function of DRG neurons, especially nociceptors, in the developing rabbit is warranted.

In adults, physical trauma is known to alter neural circuits and have long lasting effects on nociception (Costigan et al., 2009; Price and Dussor, 2014). The effect of neural injury on nociceptive primary sensory neurons is a major contributor to the development of tactile and thermal hypersensitivity and chronic pain (Hulsebosch and Coggeshall, 1981a, 1981b; Koerber et al., 1994; Ramer et al., 2012; Rodin et al., 1983; Walters, 2012; Woolf et al., 1992). After trauma, it is well-established that the afferent fibers of these nociceptors exhibit robust arborizations into the superficial and deep dorsal horn of the spinal cord (Bedi et al., 2010; Detloff et al., 2016, 2014; Krenz and Weaver, 1998; Ondarza et al., 2003; Weaver et al., 2001). Thus, it is possible that these primary afferents make inappropriate synaptic connections on dorsal horn neurons subserving non-nociceptive functions such as low threshold mechanosensation or touch that may not occur during normal development (Abraira et al., 2017; Abraira and Ginty, 2013; Boyle et al., 2019; Peirs et al., 2021). Piers et al (Peirs et al., 2021) demonstrate that spinal cord circuitry that transmits nociceptive information is dependent upon the nature or etiology of hypersensitivity or pain behaviors. Physiologically, nociceptors increase spontaneous activity and become hyperexcitable following CNS trauma (Bedi et al., 2010; Burchiel et al., 1985; Gold and Gebhart, 2010; Gracely et al., 1993; Li et al., 2018; North et al., 2019; Pitcher and Henry, 2008).

In neonates, effects of CNS trauma (like CP or brain injury) on nociception are less studied, however there is a wealth of information on the effects peripheral nerve injuries and inflammation (Fitzgerald, 1985; Himes and Tessler, 1989; Pattinson and Fitzgerald, 2004; Shortland and Fitzgerald, 1994; Torsney and Fitzgerald, 2002; Walker et al., 2003). Peripheral inflammation enhances collateral sprouting of primary nociceptive afferents and enhances nociception in adulthood (Ruda et al., 2000). We would be remiss if we did not also postulate that perinatal HI may also induce anatomical and physiological plasticity of other subtypes of primary afferent fibers or supraspinal centers involved in pain perception or pain modulation. Anoxia in the neonatal rat led to paw hypersensitivity and significantly fewer cells in the ventroposterolateral nucleus of the thalamus and somatosensory and insular cortex (Helou et al., 2021; Kumar et al., 2020, 2017).

Neuroinflammation, nociception, and pain are tightly intertwined (Grace et al., 2014). Astrocytes and microglia are activated by neural injury and release a plethora of pro-inflammatory molecules into the spinal milieu, and their activation corresponds to the development of neuropathic pain (Chhaya et al., 2018; Detloff et al., 2008; Gao et al., 2009; Hulsebosch et al., 2009; Ishikura et al., 2021; Lu et al., 2021; Xie et al., 2009; Zhang et al., 2016, 2007). After complete spinal cord injury, increased GFAP persists for at least 3 weeks in the adult rat spinal cord (Choi et al., 2021; Ducza et al., 2021; Morin-Richaud et al., 1998; Nesic et al., 2006; Xu et al., 2021). However, the time course of the GFAP response to injury is different in neonates. Unlike adults, neonatal rat pups respond to spared nerve injury with an early increase in GFAP (one day after injury), along with the same elevation after 7 days that is observed in adults (Vega-Avelaira et al., 2007). Density of GFAP fiber length and density of GFAP+ cell bodies are increased in the lumbar gray matter of motor-affected (hypertonic) HI rabbit kits at P1, seven days after HI (Drobyshevsky and Quinlan, 2017; Synowiec et al., 2019). The current study was performed in HI kits that were 4 days older (P5) and examined specific laminae/tracts involved in cutaneous sensation or nociception: (1) the dorsal column, the ascending pathway for cutaneous sensation; (2) the superficial dorsal gray matter, containing second-order projection neurons relaying nociception; (3) the ventral spinothalamic tract, carrying crude touch and pressure information; and (4) the lateral spinothalamic tract, the ascending pathway for nociception. While astrogliosis is not present in the spinal regions of interest we studied at P5 (11-14 days after HI), their reactivity closer to the time of injury could be a causal link of enhanced nociception as has been demonstrated in other models (Milligan et al., 2000). Astrocytes have been shown to exhibit different phenotypes or reactive states following similar CNS injuries that can be either destructive or pro-reparative (Escartin et al., 2021; Liddelow and Barres, 2017). Moreover, it has been reported that ischemia can induce an “A2-like” astrocyte phenotype that upregulates thrombospondins and neurotrophic factors that promote survival and repair of neurons (Gao et al., 2005; Hayakawa et al., 2014; Zador et al., 2009). Future studies could examine more closely the time course and specific regions of elevated astrocyte activation in the rabbit after prenatal HI and its relationship to enhanced nociception observed here at P5.

The stereotypical aversive paw withdrawal response to nocifensive tactile or thermal stimuli is mediated by a segmental spinal reflex and is often inferred as a proxy for cortical perception of the stimulus. It is unclear whether the results of the von Frey and Hargreaves’ tests performed here reflect hyperreflexia of nociceptive segmental spinal cord circuits (Advokat, 2002; Kauppila et al., 1998; Murphy and Zemlan, 1990; Schomburg and Steffens, 1998) or the complicated perception of neuropathic pain. It is likely these stimuli were offensive to the rabbits because paw withdrawal was typically accompanied by paw licking and/or a rapid jump away from the stimuli. Unlike in rodent experiments, there are surprisingly few reports measuring pain or nociception in experimental rabbit models, even those modeling knee osteoarthritis or intervertebral disc dislocation (Lei et al., 2017; Mans, 2020; Yoshioka et al., 1996). Composite pain scales for rabbits in a clinical environment (Banchi et al., 2020) and the Bristol Rabbit Pain Scale (Benato et al., 2021) examine and score rabbit facial expressions and other behaviors to assess pain, such as the grimace scale (Evangelista et al., 2021; Keating et al., 2012). Yet, recent reports suggest that the experimenter’s presence can have a profound effect on the animal’s behavior and subsequent score on these tests (Pinho et al., 2020). In the present study, unlike assessments of spontaneous pain using the rabbit grimace scale, these assessments identify the dermatomal location and relevant spinal cord or dorsal root ganglion level in which dysfunctional sensation occurs. This is especially useful in examining changes in primary afferent input and/or dorsal horn processing in specific spinal segments. Also, since our study examined an early postnatal age (P5) when the neonatal rabbits’ eyes are not yet open, the tests we performed were more appropriate to assess nociceptive behavior than using the grimace scale, place-preference or aversion tests. However, future work could include assessments of pain perception and an associated fear response at later postnatal ages to further characterize the perception of pain and translate these findings clinically.

Individuals with CP have disruptions in somatosensation in addition to motor dysfunction. Disruptions in somatosensory cortex connectivity (Hoon et al., 2002) and increased latency of sensory evoked potentials (Cahan et al., 1987; Kundi et al., 1989) have been well documented in individuals with CP. Innocuous tactile sensitivity actually decreases in individuals with CP while pain sensitivity is enhanced (Riquelme and Montoya, 2010). The mechanism of this is not clear, but it is possible that this is due to afferent switching after injury: in a rat model of neuropathic pain, large diameter cutaneous/somatic Aβ afferents can undergo fiber-type switching after nerve ligation and begin expressing CGRP, like nociceptive fibers (Nitzan-Luques et al., 2011). This switch in identity corresponds to a period of tactile hypersensitivity. Another possibility is that the impaired CST creates a permissive environment for afferent sprouting. When competition between CST and afferent input to the spinal cord is lost after transection of the pyramidal tracts in adult rats, afferent sprouting into the deep dorsal horn is observed (Jiang et al., 2016; Tan et al., 2012). Afferent sprouting after CST transection is accompanied by loss of rate dependent depression of the H reflex, a hallmark of CP (Tan et al., 2012). The common thread in both of these mechanisms is the remarkable plasticity of afferents, even in adulthood. This raises the possibility that aberrant nociceptive input either by an increased anatomical distribution and/or functional /physiological alterations arising during perinatal development could be corrected later in life.

Movement and motor unit firing patterns in neurotypical adults are directly affected by pain (Hodges and Tucker, 2011; Martinez-Valdes et al., 2020), perhaps through neuromodulators including serotonin (5HT) (Mesquita et al., 2020). In our study, we found that HI kits were more sensitive to mechanical and thermal stimuli than sham control kits. When comparing HI kits that were motor-affected and those that were motor-unaffected, motor-affected HI kits were less sensitive to mechanical stimuli to their hindpaws while their responses in forepaws and to thermal stimuli was unchanged. We attribute this to the greater difficulty these kits had in making movements to respond to stimuli due to their hypertonia. No kits in this study were completely incapable of movement and unable to react physically to the stimuli, but their hypertonia could have impeded their reaction. Differences between motor-affected and motor-unaffected HI kits and the relationship between nociceptive afferent feedback and motor dysfunction is the focus of ongoing studies. This study represents the first evidence of altered nociceptive afferents in a model of CP.

Each individual’s pain perception includes complex psychosocial components that were not the focus of this study. In situations where both nociception and the affective experience of pain can be altered, such as after HI, it is critical to accurately identify and classify aberrant sensory behavior. Here, we utilized methods to assess and quantify shifts in sensitivity to mechanical and thermal stimuli that are sensitive, reliable, and validated in models of central and peripheral neuropathic pain across species. We found that prenatal HI heightened nociception at the level of spinal nociceptors and reflexive responses to mechanical and thermal stimuli. It could be that CP-causative perinatal injuries contribute to other aspects of the pain experience in individuals with CP, including changes to mood and cognition, in addition to altered nociception. Thus, treatment options for chronic pain in individuals with CP should be explored with greater specificity, considering that the pathophysiology of their pain could be distinct from that of chronic pain derived from other etiologies. Novel targets for fully alleviating CP-induced chronic pain should be explored specifically within the context of treating individuals with CP or those who have survived perinatal physical trauma.

## Conclusions

The current study demonstrates enhanced nociceptive behavior and nociceptive primary afferent distribution in neonatal rabbits after prenatal HI injury. Identification of how prenatal physical stressors impact nociceptors could be the first step to preventing aberrant nociception and ultimately chronic pain in those at risk.

## Author contributions

EJ Reedich contributed to data curation, formal analysis, investigation, methodology, supervision, validation and writing, editing and reviewing the manuscript. LT Genry contributed to data curation, formal analysis, investigation, methodology, supervision, validation, and reviewing and editing the manuscript. MA Singer contributed to data curation, formal analysis, investigation, methodology, and validation. CF Cavarsan contributed to formal analysis, investigation, methodology, and validation. E Mena Avila contributed to data curation, formal analysis, investigation, methodology, validation, and reviewing the manuscript. DM Boudreau contributed to data curation, formal analysis, investigation, methodology, validation, and reviewing the manuscript. MC Brennan contributed to data curation, formal analysis, investigation, methodology, validation, and reviewing the manuscript. AM Garrett contributed to data curation, formal analysis, investigation, methodology, validation, and reviewing the manuscript. L Dowaliby contributed to investigation, methodology and supervision of this project. MR Detloff contributed to contributed to conceptualization, data curation, funding acquisition, project administration, resources, supervision, and writing, editing and review of the manuscript. KA Quinlan contributed to conceptualization, data curation, funding acquisition, project administration, resources, supervision, and writing, editing and review of the manuscript.

## Abbreviations

(5HT): Serotonin
(CGRP): calcitonin gene-related peptide
(CNS): central nervous system
(CP): cerebral palsy
(DAPI): Diamidino-2-Phenylindole
(DC): dorsal column
(DRG): dorsal root ganglion
(GFAP): glial fibrillary acidic protein
(GMFCS): Gross Motor Function Classification System
(HI): hypoxia-ischemia
(IB4): isolectin B4
(IHC): immunohistochemistry
(LST): lateral spinothalamic tract
(NHS): normal horse serum
(NSAIDs): nonsteroidal anti-inflammatory drugs
(P): postnatal day
(PBS): phosphate buffered saline
(PFA): paraformaldehyde
(ROIs): regions of interest
(SDGM): superficial dorsal gray matter
(VST): ventral spinothalamic tract

## Other Acknowledgements

This project was supported by funding from NIH NINDS including NS097880 to MRD and NS104436 to KQ. MCB and DMB received a grant from the University of Rhode Island Undergraduate Research & Innovation. The funders of this project had no role in study design, the collection, analysis and interpretation of data, in writing the report, or in the decision to publish these results.

## Data Accessibility

The data that support the findings of this study are available from the corresponding author upon reasonable request.

---

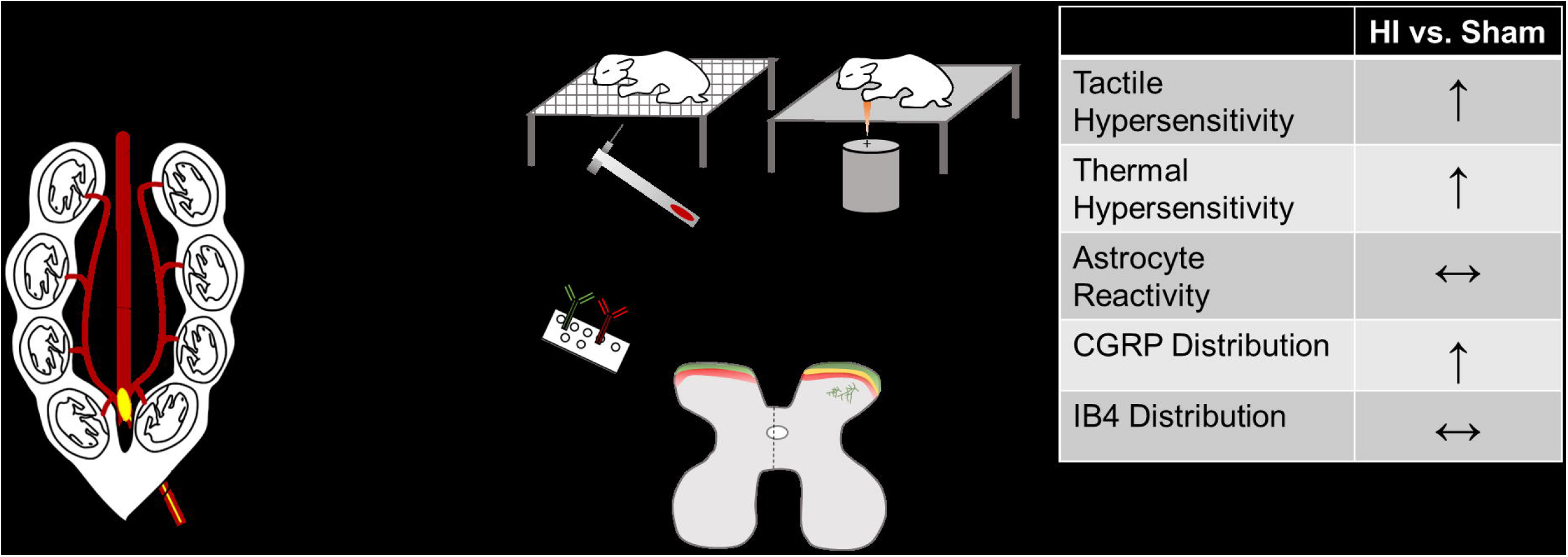

